# High defense system expression broadens protection range at the cost of increased autoimmunity

**DOI:** 10.1101/2023.11.30.569366

**Authors:** Nitzan Aframian, Shira Omer Bendori, Tal Hen, Polina Guler, Avigdor Eldar

## Abstract

The evolutionary arms race between bacteria and their phage viruses has given rise to elaborate anti-phage defense mechanisms. Major advances have been made in revealing the molecular details underlying diverse defense systems, but general principles and constraints are largely unkown. Defense systems are often tested against a diverse set of phages, revealing widely varying protection ranges. While these disparities are usually attributed to differences in mechanism, here we show that increasing expression of defense systems can greatly enhance their protection range. This holds true for disparate mechanisms, and is achieved by circumventing phage strategies for overcoming bacterial defense. However, increased defense system expression comes with a heavy cost of autoimmunity. Therefore, the expression level of defense systems controls a tradeoff between protection range on the one hand and autoimmunity on the other. We discuss how this tradeoff may drive the regulation of defense systems expression and the acquisition of multiple systems within the same genome.

## Introduction

Every known life form is plagued by viruses, and bacteria are no different. Accordingly, it is increasingly appreciated that bacteria employ a diverse set defense system against their viruses, called phages^1–3^. A basic step often performed in the characterization of newly found systems, is to outline their protection range by testing their ability to defend against a variety of phages. While some systems have been shown to protect bacteria against a broad set of phages^2,4–6^, others seem to confer a far narrower protection range, defending only against particular phages^7,8^.

Great progress has been made in understanding how the underlying molecular mechanisms of defense affect the protection range afforded by different systems. For instance, different homologs of the CapRel defense system confer protection against different phages^9^. It has been shown that this may be explained by phage-specific variations in particular residues of the major capsid protein, which acts as the trigger of this system. Accordingly, residues of CapRel thought to directly bind the major capsid protein are highly variable, suggesting ‘Red Queen’ evolutionary dynamics^9^. In contrast, different homologs of the Avs system were shown to confer extremely broad protection range due to recognition of widely conserved structural patterns of phage components that share very low sequence identity^6^.While mechanisms of defense are rapidly being elucidated, other factors affecting the protection range of different systems remain largely unexplored. Anecdotally, mechanism-independent factors such as temperature^7,10^ and expression level^11^ have been found to affect the protection range of different defense systems. However, systematic investigation of such factors, particularly factors that bacteria can control, is lacking.

Here, we explore the consequences of defense system expression levels. We find that increasing the expression of diverse defense systems can substantially enhance their protection range. Our results agree with the hypothesis that this is a consequence of overcoming different phage strategies to counter defense. However, we also find that increased defense system expression comes at an increased cost to bacteria. We show that this cost does not result from the energetic burden of expression but is rather due to the activity of defense systems, and is therefore a form of autoimmunity. The tradeoff between protection range and expression-associated cost may provide a potential explanation for the low native expression of different systems, density-dependent expression regulation^12–14^, and the acquisition of multiple specialized defense systems within the same genome^1^.

## Results

### Increased expression of the *spbK* defense system extends its protection range

To test the hypothesis that increased defense system expression leads to increased protection range (Fig. 1A), we began by focusing on the newly discovered defense system, SpbK. SPbK is a single-gene abortive infection defense system that naturally resides on the ICE*bs*1integrative and conjugative element, native to lab strains of *Bacillus subtilis*, which also host the SPβ prophage^8^. This system was shown to confer protection against phage SPβ by recognizing the YonE phage protein, and initiating growth arrest in response^8^. At native *spbK* expression levels, no defense was found against several other phages tested, indicating a relatively narrow range of protection. To test the effect of *spbK* expression levels, we constructed a strain with *spbK* under its native promoter (*spbK*^wt^), and a strain with *spbK* under an IPTG-inducible promoter (*spbK*^ind^). To focus specifically on the effect of *spbK* both these strains harbor *spbK* alone without the full ICE*bs*1. First, we used RT-PCR to better understand the relative expression levels of native and induced *spbK*. We found expression of *spbK*^wt^ to be relatively low, corresponding to induction with less than 5 µM IPTG (Figure S1A). This is comparable to *spbK* expression levels when residing within ICE*bs*1.

**Figure 1:**
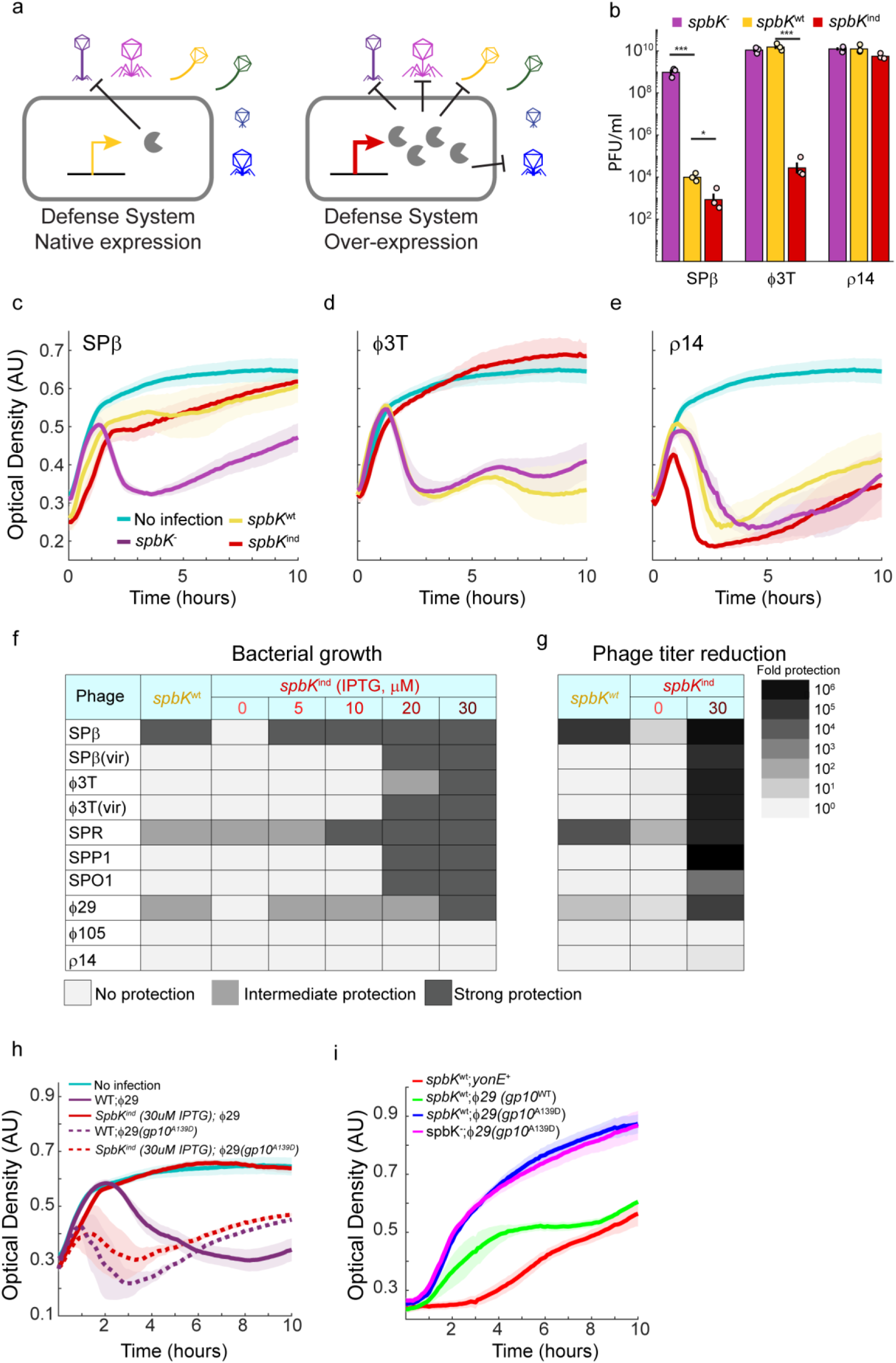
Over expression of the *spbK* defense system extends its protection range. (a) Illustration of the extension hypothesis. (b-e) PFU (b) and growth curves (d-e) after infection of phages Spβ (b,c), ϕ3T (b,d) and ρ14 (b,e). Phages infected either a defenseless background strain (*spbK*^-^, purple), a strain encoding for *spbK* under its native promoter (*spbK*^wt^ strain, yellow) or a strain overexpressing SpbK (*spbK*^ind^ strain with 30μM IPTG, dark red). In (b) the geometric means and error bars represent the s.e.m. of the log are shown. Empty circles represent n = 3 biological replicates; * marks p<0.05, *** marks p<0.0005. In (d-e). mean OD of n=3 biological replicates and and s.e.m. (lighter shade) are shown. Growth of non-infected cells is also shown in cyan. (f,g) Level of protection inferred from (f) growth curve and (g) PFU for 10 designated phages infecting either *spbK*^wt^ or *spbK*^ind^ with different levels of IPTG (See also Figure S2and S3). Color code for protection levels is explained in the legends. (h) Comparison of infection curves of phages ϕ29 (solid line) and its escaper mutant ϕ29(gp10^A139D^) (dashed line) upon infection of wild-type (purple) and *spbK*^ind^ (30μM IPTG, deep red) strains. (i) Growth curves of cells coexpressing the natively expressed *spbK* gene and either of three xylose-induced portal proteins; *yonE* (red) from phage SPβ, *gp10* from phage ϕ29 (green) or *gp10^A^*^139^*^D^* escaper mutation (blue). The latter strain is also shown with no *spbK* gene coexpression as a control (magenta).

Next, we tested the protection range of *spbK* at different expression levels against a diverse set of phages by tracking bacterial growth during infection as well as phage propagation 2 hrs post infection (Methods; Fig 1B-G, Figure S2, S3). To this end we used a diverse set of 10 phages and phage mutants that differ in their lifestyle and belong to different morphological families. All infection experiments were performed on bacteria grown to mid-log phase in liquid lysogeny broth (LB) at a multiplicity of infection (MOI) of 0.1. In terms of growth curves, we differentiated no reduction in OD (strong protection), and a diminished or delayed reduction in OD (weak protection) (Fig. 1C-F; Figure S3). At native expression levels, we found that *spbK* strongly protects against phages SPβ and SPR with observed ∼5 and ∼4 orders of magnitude reduction in phage titer respectively, and no inhibition of bacterial growth (Fig 1B-G). Apart from these phages, which both belong to the SPβ-like family^15–17^, natively expressed *spbK* offered only weak protection against phage ϕ29 (a 10-fold reduction in phage production that did not prevent bacterial population collapse). Increasing expression led to protection against some additional phages but not others. For example, overexpression of *spbK* conferred potent protection against phage ϕ3T – phage propagation was reduced by ∼5 orders of magnitude, and bacterial growth curves during infection did not observably deviate from infection-free growth curves (Fig. 1B, D; Figure S2, S3). In contrast, phage ϕ105 remained unaffected even when *spbK* was overexpressed – no discernable defensive effect of native nor overexpressed *spbK* was found in terms of bacterial growth or phage propagation (Fig. 1B, E; Figure S2, S3).

Overall, protection range dramatically increased with increased *spbK* expression – at 30 µM IPTG, which corresponds to a ∼20-fold increase compared to native expression (Figure S1A), SpbK provided potent protection against all phages tested, with the exclusion of ϕ105 and ρ14 which both belong to the ϕ105-like family (Fig. 1F, G). In terms of growth curves, we found that some offered only weak protection, while others allowed for completely uninterrupted population growth. We also found that when *spbK* was overexpressed, phage propagation was reduced by more than 3 orders of magnitude for each of these 8 phages. Notably, overexpressed, but not natively expressed *spbK*, could not protect against a virulent mutant of SPβ deleted in its master repressor^18,19^. This difference between WT SPβ and its virulent mutant may stem from a more rapid infection cycle of the mutant^20^.

### SPbK can be triggered by divergent portal proteins

Interestingly, SPP1, ϕ29 and SPO1 are among the phages strongly inhibited by overexpressed *spbK*, but they do not code for a protein with high sequence identity to the SPbK-triggering YonE protein of SPβ. YonE was previously proposed to function as a portal protein based on homology search and its vicinity to a putative terminase^8^. Portal proteins of diverse phages tend to have a similar structure despite little sequence identity, and it was recently discovered that some Avs systems can recognize highly divergent portal proteins ^6^. We therefore hypothesized that SPbK can similarly recognize diverse portal proteins, even when those share little sequence identity.

To test this, we focused on the portal protein of phage ϕ29, Gp10, which has been characterized structurally and functionally ^21–23^, and shares a low (12.5%) amino acid sequence identity with YonE. We infected bacteria expressing high levels of SPbK that are sufficient to hinder ϕ29 propagation in liquid LB, and performed plaque assay on defenseless bacteria after 2 hrs. We then tested phage isolates from several plaques for their ability to infect bacteria that express high levels of *spbK* by tracking growth curves, confirming 8 escaper mutants (Fig. 1H). We sequenced *gp10* from every escaper mutant and found that they all harbored the same point missense A139D mutation, which is located at an exposed residue that faces outwards from the portal ring, and is predicted to be highly variable (methods)^23,24^. The fact that all isolates harbored the same mutation may have resulted from a single mutant taking over the population during infection in the well-mixed liquid culture, or from multiple parallel mutations.

Notably, escaper mutants caused an earlier population collapse when infecting the *spbK*^-^, defenseless, stain compared to their ancestor, followed by population recovery (Fig. 1H). The ability of these mutants to escape defense may result from altered infection dynamics that may not directly reflect on the role of Gp10 as a trigger for SpbK. To validate that SpbK can be triggered by the portal Gp10 protein of ϕ29, we co-expressed it in bacteria along with natively expressed *spbK* (Fig. 1I). Highly expressed *gp10* resulted in growth inhibition in the presence of SpbK, but not in its absence. Though substantial, the toxicity exhibited by co-expression of *spbK* and *gp10* was weaker than that of *spbK* and *yonE*. This is in line with the observation that SpbK strongly protects against ϕ29 only when overexpressed. In contrast, co-expression of *spbK* along with the mutant *gp10* found in ϕ29 escapers did not result in growth inhibition, pointing to a reduced ability to trigger SpbK. Overall, these results indicate that during phage infection SpbK can potentially be triggered by diverse portal proteins to provide potent defense. This potential is realized at high levels of expression, facilitating a broadening of protection range.

### Extension of protection range upon overexpression is found in three additional defense systems; Lamassu, Gabija and Septu

We wondered if this pattern of protection range increasing along with expression could be generalized to other defense systems. To test this, we examined three additional defense systems from different *Bacillus* species – Gabija, Lamassu, and Septu^2^, each of which was cloned under its native promoter or under an inducible promoter (either IPTG or xylose induction) into *Bacillus subtilis*. We tested the protection capability of each of these systems against the same set of 10 phages and lytic phage mutants. We found that overexpression increased protection range for all systems as measured by growth curves, attesting to the generality of the relationship between expression level and protection range (Fig 2A, Figure S4-S6).

**Figure 2:**
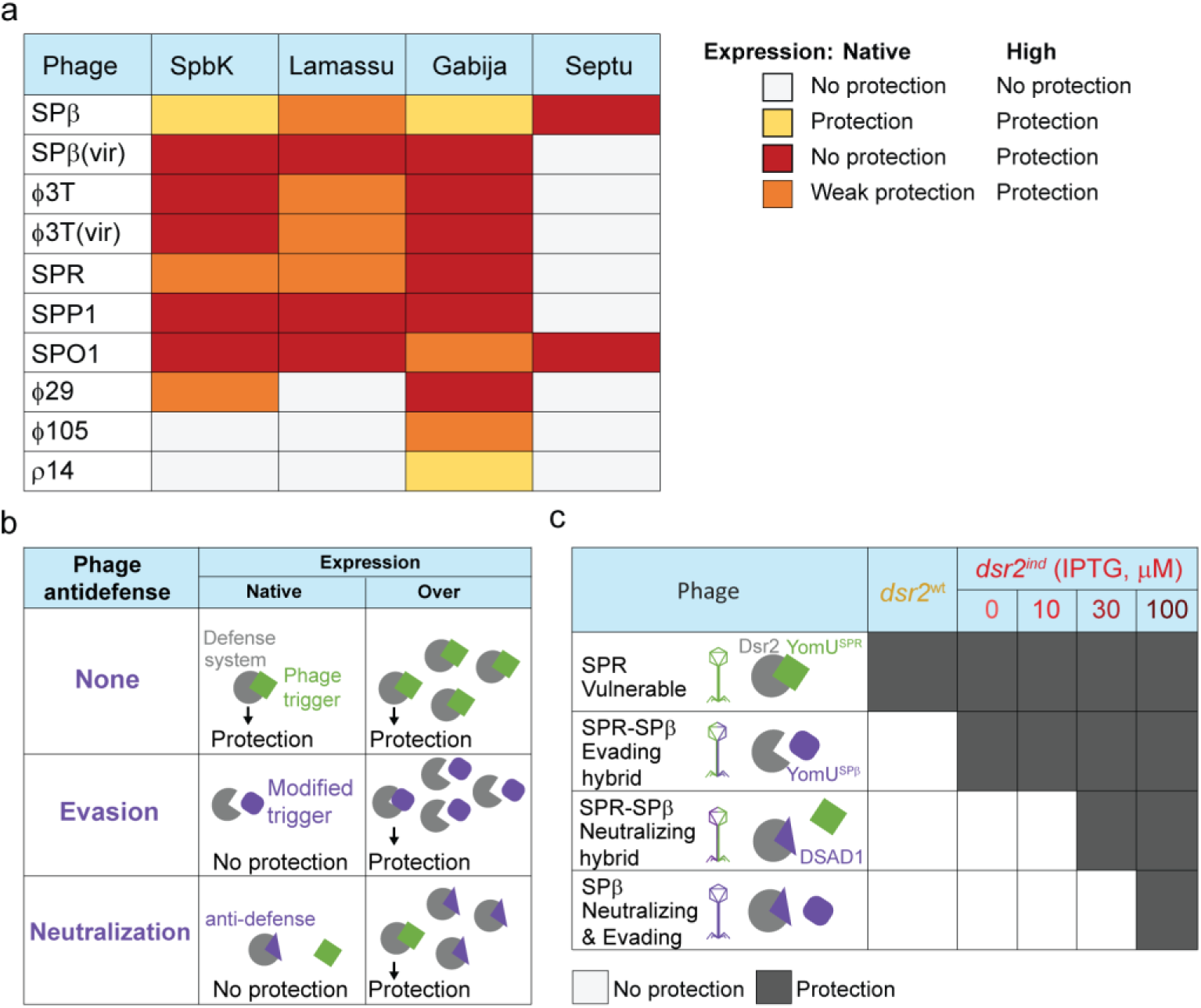
Extended protection range upon overexpression occurs in multiple defense systems and depends on overcoming different counter-defense strategies. (a) A table summarizing protection afforded by defense systems as native expression and when overexpressed, as inferred by growth curves (See also Figure S3-S6). Color code for protection levels is explained in the legends. (b) Illustration of a hypothesis for the mechanism underlying extended expression range. Increased protection overwhelms counter defense strategies of evasion and neutralization. (c) Table summarizing protection afforded by the DSR2 system at different expression levels against phages SPR, SPβ, and their hybrids. Results are based on efficiency of plating (EOP; See also Figure S7).

### Overexpression of the DSR2 defense system allows it to overcome different strategies employed by phages to circumvent defense

Next, we wanted to examine the underlying cause for the observed increase in protection range. We hypothesized that increased expression may overwhelm different strategies phages employ to circumvent bacterial defense (Fig. 2B). From a broad perspective, two strategies phages employ to overcome bacterial defense are evasion and neutralization. Evasion may be achieved, for example, by changing the phage component that acts as the trigger of defense^9,25^. Alternatively, phages can actively neutralize bacterial defense by encoding an anti-defense mechanism^11,26,27^. We reasoned that expression levels of defense systems may be crucial for the success of both these general strategies. For the evasion strategy, a lower affinity to the trigger of defense may be compensated for by a higher concentration of the defense system itself, potentially allowing bacteria to maintain rapid phage detection and effective defense activation. For the neuralization strategy, defense systems may be expressed at a level which overwhelms phage-encoded antidefense mechanisms.

Initial support for this mechanistic explanation can be found in the pattern of Gabija defense against phage ϕ3T^27^. Natively expressed Gabija defends against phage SPβ, but not against the closely related phage ϕ3T. It was recently found that ϕ3T encodes an anti-defense protein, Gad1, that blocks Gabija activity, explaining this difference^27,28^. However, here we observed that when overexpressed, Gabija does confer potent protection from phage ϕ3T, indicating that Gabija defense overwhelms phage anti-defense in this case.

To test our hypothesis more directly, we shifted our focus to a system for which we can test the interaction between expression range and mechanisms of anti-defense and evasion. It was recently discovered that a DSR2 system from *Bacillus subtilis* provides defense by initiating NAD+ depletion upon recognition of a phage tail-tube protein^29^. It was shown that phage SPR propagation is inhibited by DSR2 while the closely related phage SPβ is not. Two features explain the failure of DSR2 to defend against SPβ – first, the tail-tube protein of SPβ has lower affinity to DSR2. Second, SPβ encodes for an anti-defense protein, Dsad1, which inhibits DSR2 through direct competitive binding^29^.

In the process of deciphering the mechanism of DSR2 defense, several SPR-SPβ phage hybrids were made. We used two such hybrids for our purposes – one hybrid which possesses the tail-tube protein of SPβ but does not code for Dsad1 (hybrid J76; an ‘evading’ hybrid), and another which possesses the tail-tube protein of SPR, but additionally encodes for Dsad1 (hybrid J78; a ‘neutralizing’ hybrid) (Fig 2C). We tested the efficiency with which SPR, SPβ, and these two hybrids could form plaques on a lawn of bacteria expressing DSR2 at different levels (efficiency of plating, EOP).

As expected, under native expression, DSR2 could only defend against phage SPR. We next cloned the *dsr2* gene under IPTG-induction and found that protection range increased as we increased *dsr2* expression levels (Fig. 2C). First, we find that leaky expression from the inducible *dsr2* system was already higher than that of the native system and defended against the ‘evading’ hybrid even when no IPTG was added. At a higher expression level (30 μM IPTG), DSR2 defended against the ‘neutralizing’ hybrid as well, and finally at yet higher expression (100 μM IPTG), defense was observed even against phage SPβ, which utilizes both strategies (Fig. 2C; Supplementary Figure 7).

In total, these results show that the expansion of protection range with expression level is widespread and that it may generally be explained by overcoming different phage strategies of evasion and anti-defense.

### Sufficient increase of defense system expression leads to reduced growth

An increased range of protection could potentially be highly beneficial to bacteria, and increasing the strength of a promoter is a relatively simple evolutionary adaptation. However, our results indicate that native expression levels of defense systems are far from those maximizing protection range (Fig 1G, H; Fig. 2A, C). We reasoned that this may potentially be explained by a substantial cost associated with increased expression levels. To explore this, we first returned to *spbK* to test the expression-dependent cost of this system (Fig. 3A, B). Using the IPTG-dependent, s*pbK*^ind^ strain, we found that at high IPTG levels (>=50μM), that are beyond the range used to examine protection (Fig. 1A-H), bacteria did not grow at all or exhibited substantially delayed growth (Fig. 3A). Bacteria showed near normal growth at lower IPTG levels (<=30μM) when examined in monoculture. To better identify smaller growth defects, we used a coculture assay to compete differentially fluorescently labeled wild-type strain and a strain encoding the inducible *spbK* at different IPTG levels (methods). We find that *spbK*-expressing cells showed a sharp reduction in competitive fitness at induction levels of 30μM (P=2*10^-4, two-tailed paired t-test; Fig. 3C).

**Figure 3:**
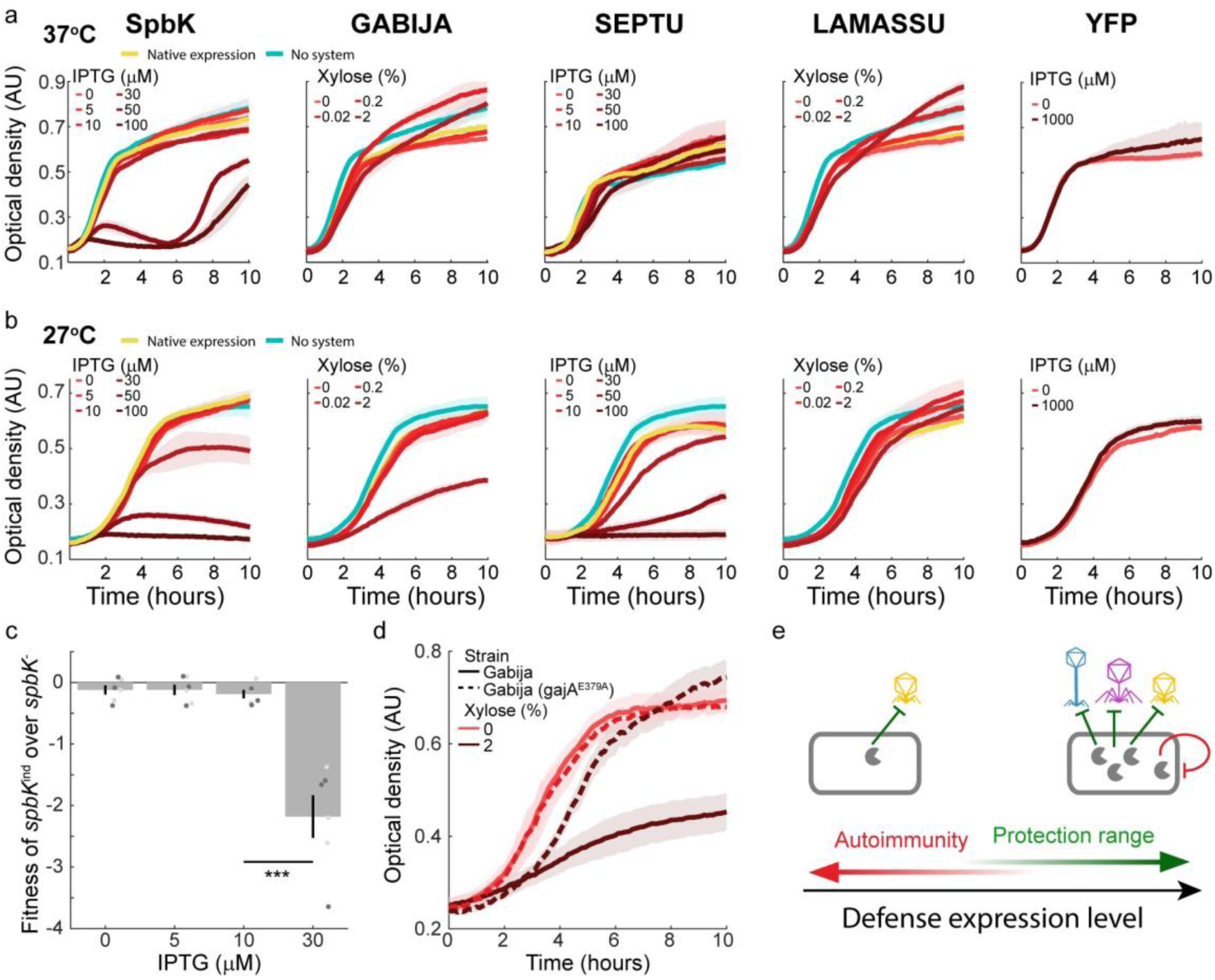
Defense overexpression leads to autoimmunity in multiple defense systems. (a, b) Growth curves of bacteria expressing defense systems or a yellow fluorescent protein (YFP) at different levels, at 37° (a) or 27° (b). Mean OD of n=3 biological replicates and and s.e.m. (lighter shade) are shown. (c) Fitness of *spbK*^ind^ over *spbK^-^* at different expression levels. Mean and s.e.m are shown. Individual points represent n=6 biological repeats. Light and dark points represent different fluorescent colors used (methods). * marks p<0.05. (d) Growth curves of strains expressing xylose induce Gabija (wild type sequence, solid lines) or a Gabija mutant (gajA^E379A^; dashed lines) in the absence (light red) or presence (dark red) of xylose. Mean OD of n=3 biological replicates and and s.e.m. (lighter shade) are shown. (e) Illustration of the expression-controlled tradeoff between the benefit of protection range and the cost of autoimmunity. Green and red arrows represent opposing selection forces towards high or low expression, respectively.

We next asked whether other defense systems exhibit an expression-dependent cost as well. By tracking growth curves in monocultures at different expression levels, we found that Septu, but not Gabija and Lamassu showed an observable growth inhibition at high expression levels (Fig 3A). In addition to expression-level, other factors may also affect the activity and cost of defense systems. Previous reports have indicated that temperature may affect the activity of different defense systems^7,10^. We found that decreasing temperatures from 37° to 27° increased the cost of SPbK, Gabija and Septu overexpression (Fig 3B): growth inhibition by SPbK was observed at lower induction levels, and Gabija and Septu showed greater inhibition for the same induction levels. Lamassu did not incur an observable cost at either temperature (Fig. 3C). Other conditions or more sensitive assays may reveal other costs of different systems. Notably, it was found that a type II Lamassu system becomes costly to cells in the presence of some plasmids^30^, suggesting that Lamassu systems may be costly under conditions other than those we tested.

### Growth inhibition at high expression levels is due to autoimmunity

The cost associated with increased expression could result from the activity of defense systems, or from alternative sources such as metabolic burden. To test the possibility of cost due to metabolic burden, we first expressed a yellow fluorescent protein (YFP) under an IPTG-inducible promoter (Fig. 3A, B). We found that YFP expression had no observable effect on bacterial growth, even at saturating inducer levels (Figure S1B). For a more direct control and to further rule out other possible sources of cost, we introduced a point missense E379A mutation in the endonuclease domain protein of the Gabija system (*gajA*), which has previously been shown to abrogate defense^31^. We found that in contrast to the WT system, the overexpressed mutant Gabija did not impose an observable cost at 27° (Fig. 3D). This suggests that the cost of the system is a direct side-effect of its defensive function, which constitutes a form of autoimmunity.

In total, our results indicate that for diverse defense systems, increasing expression levels comes with a tradeoff – on the one hand a wider protection range is achieved by overcoming different phage strategies, but on the other hand the cost of autoimmunity increases as well (Fig 3E).

## Discussion

Many studies have observed differences in protection range between different mechanisms. Here we show that increasing a systems expression can substantially alter its protection range – defense systems that under native expression afford a modest range of protection may hold the potential for a much broader protection range under increased expression levels. However, we show that for some systems increasing expression also comes with an increased cost, stemming from autoimmunity. This tradeoff may explain why native expression levels have not evolved to simply maximize protection range. We show that increased expression can increase protection range by circumventing different phage counter strategies against bacterial defense, including evasion and active disruption by anti-defense. Evidence to support this can also be found with the toxin-anti defense system ToxIN which successfully defends against phage T4 when carried on a multicopy plasmid, despite T4 carrying its own antitoxin^11^. In contrast, when integrated in a single copy in the bacterial chromosome, the defense-neutralizing antitoxin encoded by T4 is sufficient for defense.

The effectiveness of defense system overexpression against anti-defense mechanisms may come as a consequence of a basic asymmetry between bacteria and phages – while defense systems can be expressed prior to infection, expression of phage-encoded anti-defense mechanisms often begins only once infections commence^7,11,27,32–34^. Indeed, anti-defense mechanisms such as anti-CRISPRs have been shown to be expressed very highly and very early during infection, which may assist phages to compensate for the initial disparity in favor of the pre-expressed defense^26,35^. An alternative strategy to negate this asymmetry between defense and anti-defense is by packaging anti-defense proteins into mature virions such that these are injected into a host along with the phage genome^11,36^.

Importantly, here we performed all experiments under a low MOI (0.1), but MOI may be an additional variable affecting protection range. It has been shown that some anti-CRISPRs are only effective at high MOIs^32,37^. This is the result of a cooperative effect in which the culminative immunosuppressive effect of anti-CRISPRs expressed by phages serially infecting single bacteria eventually leads to effective CRISPR inhibition and phage propagation. A similar MOI-dependent effect is likely to occur in other anti-defense mechanisms.

Several studies have previously examined the cost of defense systems, with varying results. For instance, while the λ-phage-encoded RexAB system was reported to have no observable cost in terms of growth rate or competition experiments^38^, the anti-T4 Lit system was reported to incur a major cost when introduced on a high copy plasmid into *E. coli* cells^39^. Restriction-modification systems have been observed to occasionally cleave the bacterial chromosome, but this was only shown to be costly in DNA-repair deficient mutants^40^. Additionally, it has been hypothesized that the evolutionary pattern of frequent gain and loss of defense systems may be explained by a major cost they incur^41–43^. Here we examined the cost of defense systems as a function of expression level, showing that several divergent systems incur a major expression dependent cost stemming from autoimmunity.

In contrast to other systems, Lamassu did not exhibit an observable cost on growth rate even at high expression levels. As noted above, the cost of Lamassu may be more pronounced in the presence of plasmids^30^. More generally, the cost of defense systems may be condition-dependent. A striking example of condition-dependent cost is that for all defense systems for which cost was observed, it was exasperated under lower temperatures. It would be intriguing to pursue the source of this phenomenon in future studies. More broadly, there are other potential aspects of the cost of defense systems which we have not examined here. For instance, there could be an infection-dependent cost of defense systems, as many systems target core cellular processes upon activation^44–47^, but not all phage infections are lethal. A prime example is infection by temperate phages which could result in lysogeny.

The expression-controlled tradeoff between protection range and autoimmunity may have several implications. Firstly, it suggests that regulation of expression is important – high expression is very beneficial when infections are likely, but detrimental when they are unlikely. It has been shown that the expression of some CRIPSPR systems is controlled in a density-dependent manner, allowing for increased expression at high bacterial densities when phages are more likely to spread throughout the population^12–14^. Additionally, some systems have been reported to be induced upon phage infection^48–53^, though such late expression may fail to defend against rapidly replicating phages. These may be seen as a possible solution to the tradeoff we describe, and we envision that other forms of expression regulation may be revealed in the future. A wholly different, but not mutually exclusive, solution to the tradeoff we observed is possessing several defense systems, each specialized against a different set of phages. This is also in agreement with some patterns of interactions between phages and defense systems observed in natural populations^49,54^. An assortment of multiple, lowly expressed, defense systems may achieve a broad protection range, without the major cost associated with a single, highly expressed, system. Therefore, the tradeoff we describe here may be a potential driver of the frequent co-residence of multiple defense systems within the same genome^1^, even before considering possible synergistic interactions or layered activation of anti-phage defense^42,49,55^.

## Methods

### Strain construction

All bacterial strains, plasmids and primers used in this study are listed in Supplementary Tables 1–3. To construct the new *B. subtilis* strains, standard transformation protocols were used for genomic integration and plasmid transformation^56^. All stains were constructed on the background of the PY79 *B. subtilis* lab strain. An exception are strains harboring the Septu system, which were constructed on the background of the *B. subtilis* BEST7003.

Defense systems used in this study consist of Gabija from *B. cereus* VD045, Lamassu from Bacillus sp. NIO-1130, and Septu from *B. weihenstephanesis* KBAB4, all discovered by Doron et al^2^. In addition, we used DSR2 from *B. subtilis* 29R (ref ^29^) and SpbK from *B. subtilis* strain 168. Sequences of all defense systems and native promoters used are listed in Table 4.

### Growth media and conditions

All experiments were performed in LB medium: 1% tryptone (Difco), 0.5% yeast extract (Difco) and 0.5% NaCl. Liquid cultures were grown with shaking at 220 r.p.m and a temperature of 37 °C. When preparing plates, medium was solidified by adding 2% agar. Antibiotics were added (when necessary) at the following concentrations: spectinomycin, 100 µg ml^−1^; chloramphenicol, 5 µg ml^−1^; kanamycin, 10 µg ml^−1^; macrolide, lincosamide and streptogramin, 3 µg ml^−1^; erythromycin + 25 µg ml^−1^ lincomycin. For the experiments that involved infection, MMB medium was used (LB supplemented with 0.1 mM of MnCl_2_ and 5 mM of MgCl_2_).

### Plaque forming assay

Samples for PFU measurements were collected from cultures centrifuged for 5 min at 4,000 r.p.m. at room temperature. Next, the supernatant was filtered using a 0.2 μm filter. Then, 100 μl of filtered supernatant at an appropriate dilution was mixed with 200 μl of the appropriate indicator strain grown to OD_600_ in MMB and left to incubate at room temperature for 5 min. Three milliliters of molten LB-0.5% agar medium (at 60 °C) supplemented with 0.1 mM of MnCl_2_ and 5 mM of MgCl_2_ was added, mixed and then quickly overlaid on LB-agar plates. Plates were then incubated at 37 °C for 1 h and then overnight at room temperature to allow plaques to form.

### Infection experiments

Strains were grown overnight in LB at 37° C with shaking at 220 r.p.m. then diluted by a factor of 1:100 into fresh MMB medium supplemented with the indicated concentration of inducers (either xylose or IPTG). On reaching OD_600_= 0.3, cultures were infected with phages at MOI = 0.1. To examine bacterial growth, OD measurements at a wavelength of 600 nm were then performed in a 96-well plate using a plate reader (VICTOR Nivo Multimode Plate Reader; PerkinElmer). To measure phage propagation, strains were grown at 37° C with shaking at 220 r.p.m. for 2 hrs subsequent to infection, at which point samples were collected for plaque assays as described above. Plaque assays were performed on an indicator PY79 strain that does not harbor defense systems.

### Isolation of ϕ29 escaper mutants

*spbK*^ind^ were grown overnight in LB at 37° C with shaking at 220 r.p.m. then diluted by a factor of 1:100 into fresh MMB medium supplemented with 30 μM of IPTG. Upon reaching OD_600_= 0.3, bacteria were infected with ϕ29 phages at MOI = 0.1. Lysates were collected 2 hrs post infection and plaque assays were performed on *spbK*-strains, as described above. Phages from 8 individual plaques were then used to infect *spbK*^ind^ and *spbK*^-^ strains, and bacterial growth curves were tracked as detailed above. Gene *gp10* of all escaper mutants were then sequenced, as well as that of the ϕ29 wt ancestor. Location and conservation score of the A139D mutated residue of Gp10 (PDB ID – 1JNB) was discerned using ConSurf^57^.

### Co-expression with spbK

Strains harboring *spbK* under control of wild type promoter and an additional xylose-induced gene as indicated were grown to OD600= 0.3, and then diluted by a factor of 1:10 into fresh LB containing the indicated xylose concentration. Strains were then placed in a plate reader and growth was tracked as described above.

### EOP on strains harboring DSR2

Strains were grown ON in LB at 37° C with shaking at 220 r.p.m. then diluted by a factor of 1:100 into fresh MMB medium supplemented with IPTG at the indicated concentration. On reaching OD_600_= 0.3, growing strains were used to perform plaque assay with different phages, as detailed above.

### Defense systems cost experiments

To examine bacterial growth during expression of defense systems, strains were grown overnight in LB at 37° C with shaking at 220 r.p.m. then diluted by a factor of 1:100 into fresh LB medium. On reaching OD_600_= 0.3, strains were diluted by a factor of 1:10 into fresh LB medium supplemented with different concentrations of the appropriate inducer (xylose or IPTG). and placed in the plate reader. OD measurements at a wavelength of 600 nm were then performed as described above at either 37° C or 27° C.

For competitions experiments, differentially fluorescent-marked strains were grown to OD_600_= 0.3, mixed at a 1:1 ratio. Co-cultures were then diluted at a factor of 1:10 and IPTG was added at the indicated concentration. After ON growth at 37° C with shaking strain frequencies were measured using flow cytometry (Beckman Coulter CytoFLEX flow cytometer). Relative fitness was calculated as the ratio of relative frequency of the strains at the end of the experiment to their initial frequency.

### RT–qPCR measurements

Strains grown ON were diluted by a factor of 1:100 into fresh LB medium containing appropriate concentrations of IPTG. Upon reaching OD_600_= 0.3 total RNA was extracted from cells using a High Pure RNA Isolation Kit (Roche). Samples at different time points were centrifuged for 5 min at 2,000 relative centrifugal force and pellets were flash-frozen in liquid nitrogen. Then, 1 µg of RNA was reverse-transcribed to complementary DNA (cDNA) using a qScript cDNA Synthesis Kit (Quanta BioSciences). RT–qPCR was performed on a QuantStudio™ 1 System (Applied Biosystems) using SYBR Green (Quanta BioSciences). The specificity of primers (listed in Supplementary Table 3) was validated using a melt curve. The efficiency of the primers used was between 95 and 105%, with calibration curves of r2 > 0.95. Thermocycling parameters were as follows: holding stage at 95 °C for 1 min and 40 cycles of 2 steps—a first step at 95 °C for 5 s and a second step at 60 °C for 30 s. Transcript levels were normalized to levels of the reference gene *rpoB*. RNA samples were stored at −80 °C and cDNA was stored at −20 °C. Nucleic acid quantification was performed by NanoDrop 2000c Spectrophotometer (Thermo Fisher Scientific). Results were analysed using the QuantStudio design & analysis software v1.5.2 by the standard ΔΔCt method.

## Acknowledgements

We thank Shaul Pollak, and members of the Eldar lab for fruitful discussions and comments on the manuscript. The Eldar lab is funded by a European Research Council grant 724805 and by Israel Science Foundation grant 2228/21. Nitzan Aframian is supported by the Israeli Academy of Science’s Adams Fellowship.

## Supplementary figures

**Supplementary Figure 1:**
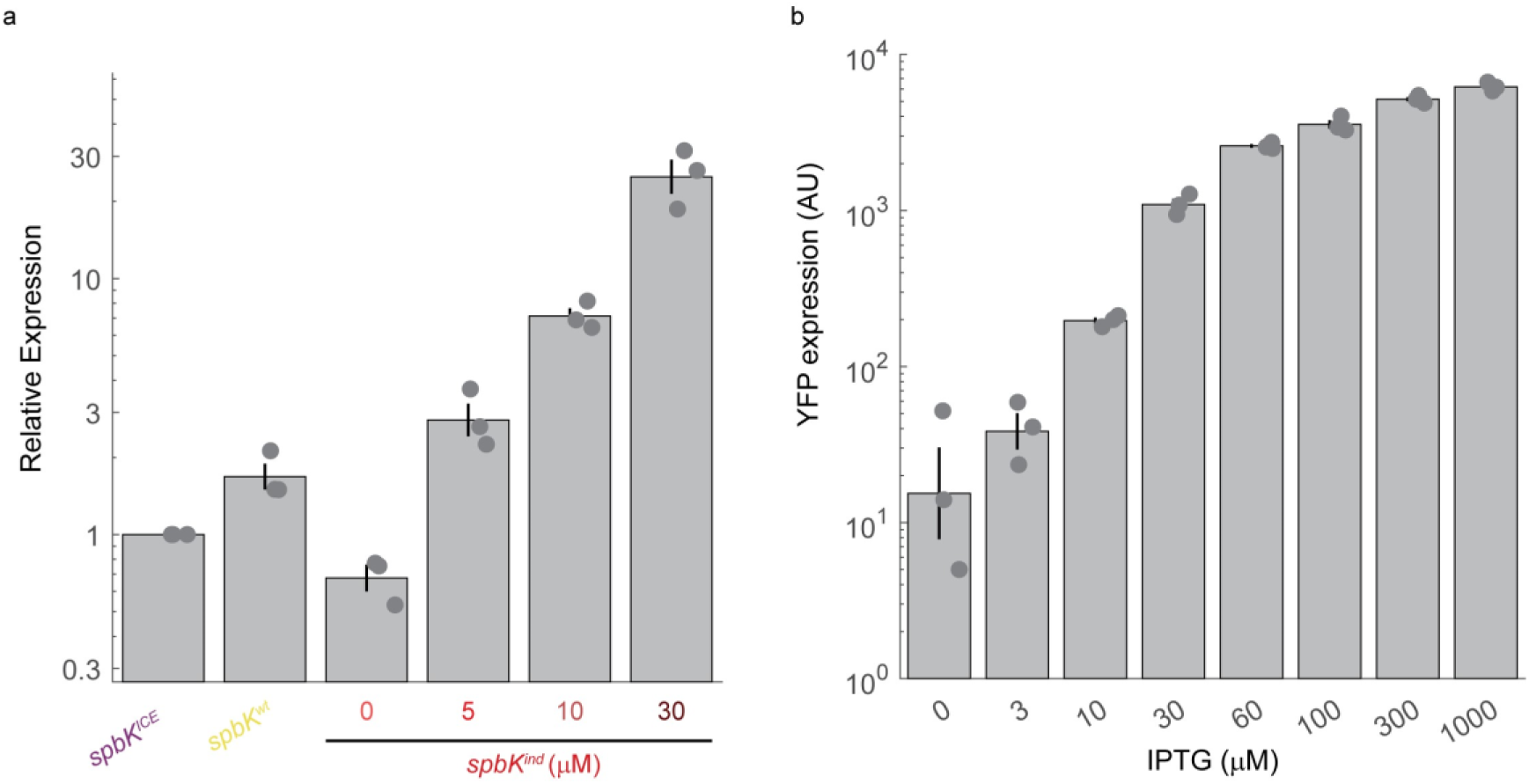
native and induced *spbK* relative expression levels. (a) Relative transcript levels of *spbK* in under different conditions, measured by RT-qPCR (methods). *spbK*^ICE^ is a strain harboring the full ICE*bs*1 element, and *spbK*^wt^ a strain harboring *spbK* alone under its native promoter. Shown are means and error bars represent s.e.m. Points represents n=3 biological repeats. (b) YFP expression under an IPTG-inducible promoter at different concentrations of IPTG. Shown are geometric means and s.e.m of the log. Individual points represent n=3 biological repeats.

**Supplementary Figure 2:**
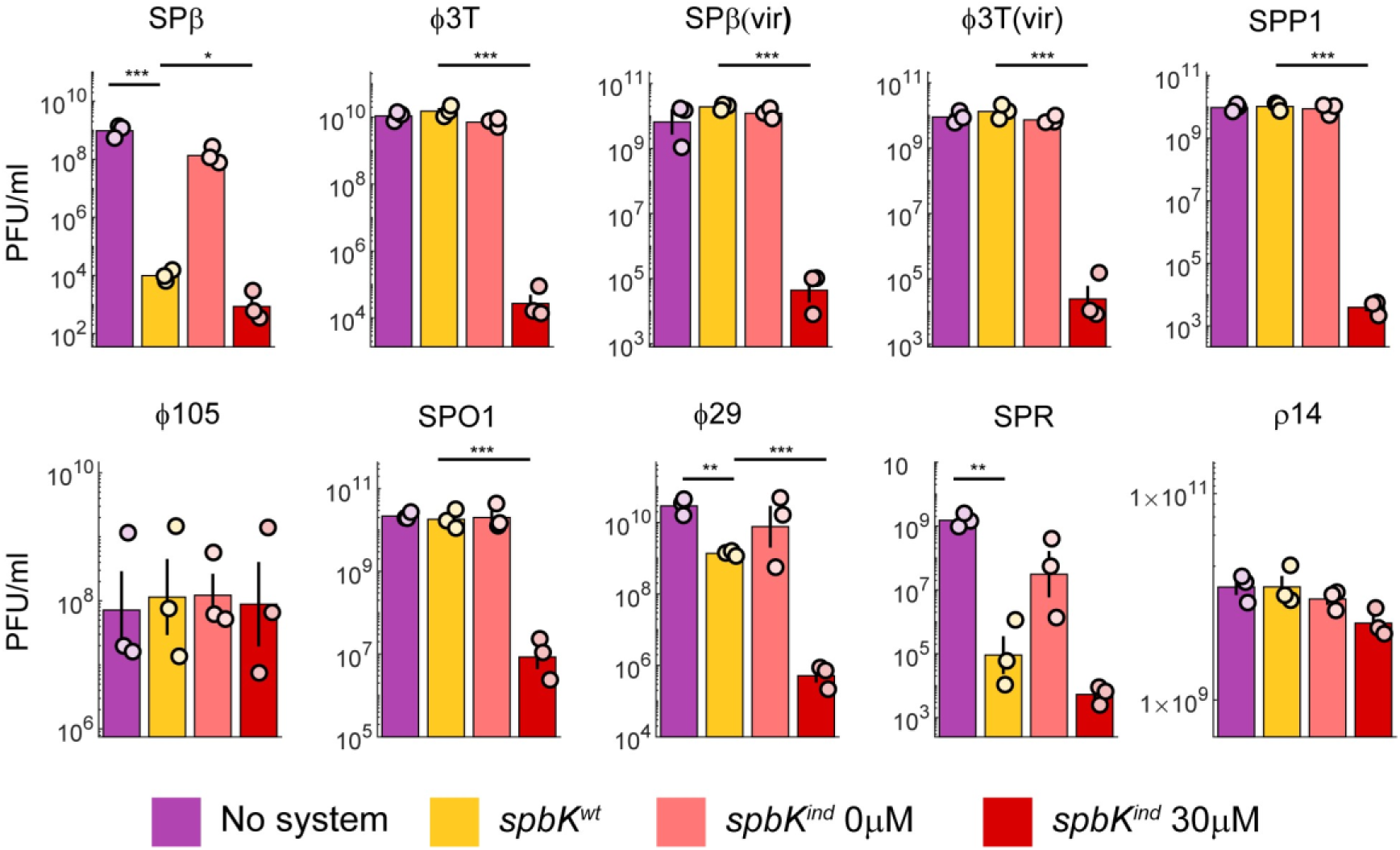
PFU/ml of different phages 2 hrs post infection of strains harboring spbK at different expression levels. Shown are geometric means and s.e.m of the log. Individual points represent n=3 biological repeats. * marks p<0.05, ** marks p<0.005, and *** marks p<0.0005.

**Supplementary Figure 3:**
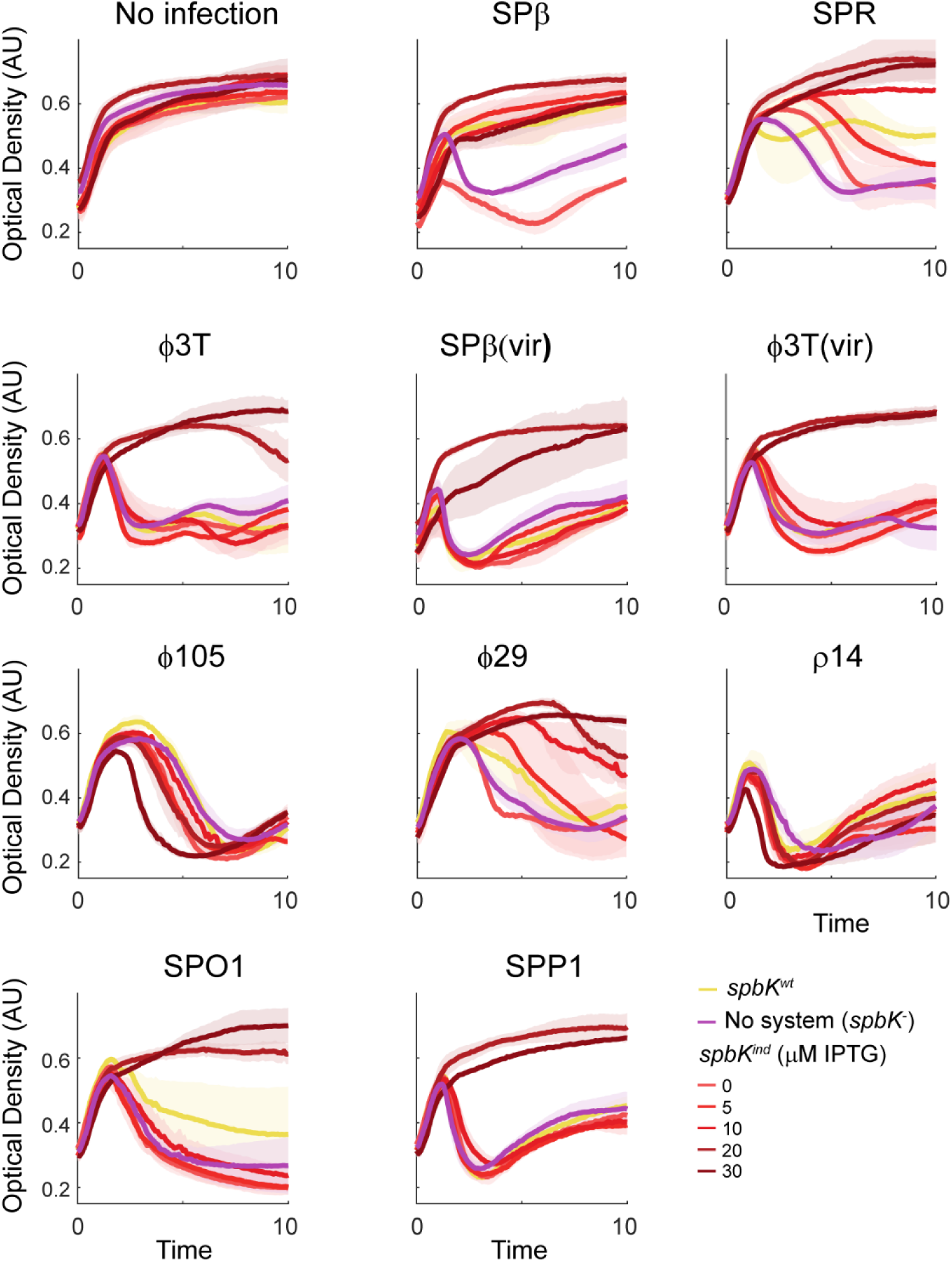
Growth curves of bacterial strains expressing *spbK* at different expression levels, infected by different phages. Mean OD of n=3 biological repeats and s.e.m (lighter shade) are shown.

**Supplementary Figure 4:**
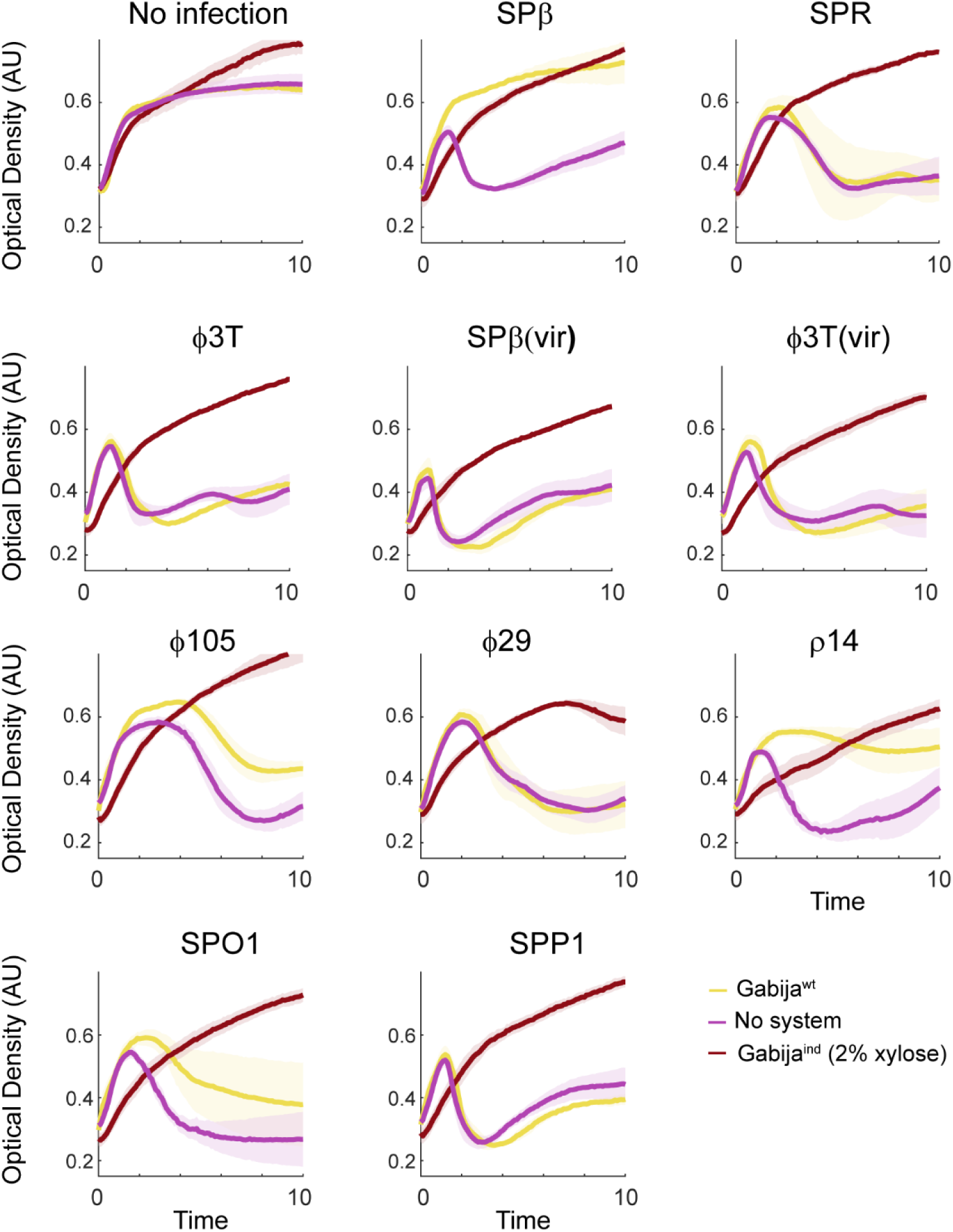
Growth curves of bacterial strains expressing Gabija at different expression levels, infected by different phages. Mean OD of n=3 biological repeats and s.e.m (lighter shade) are shown.

**Supplementary Figure 5:**
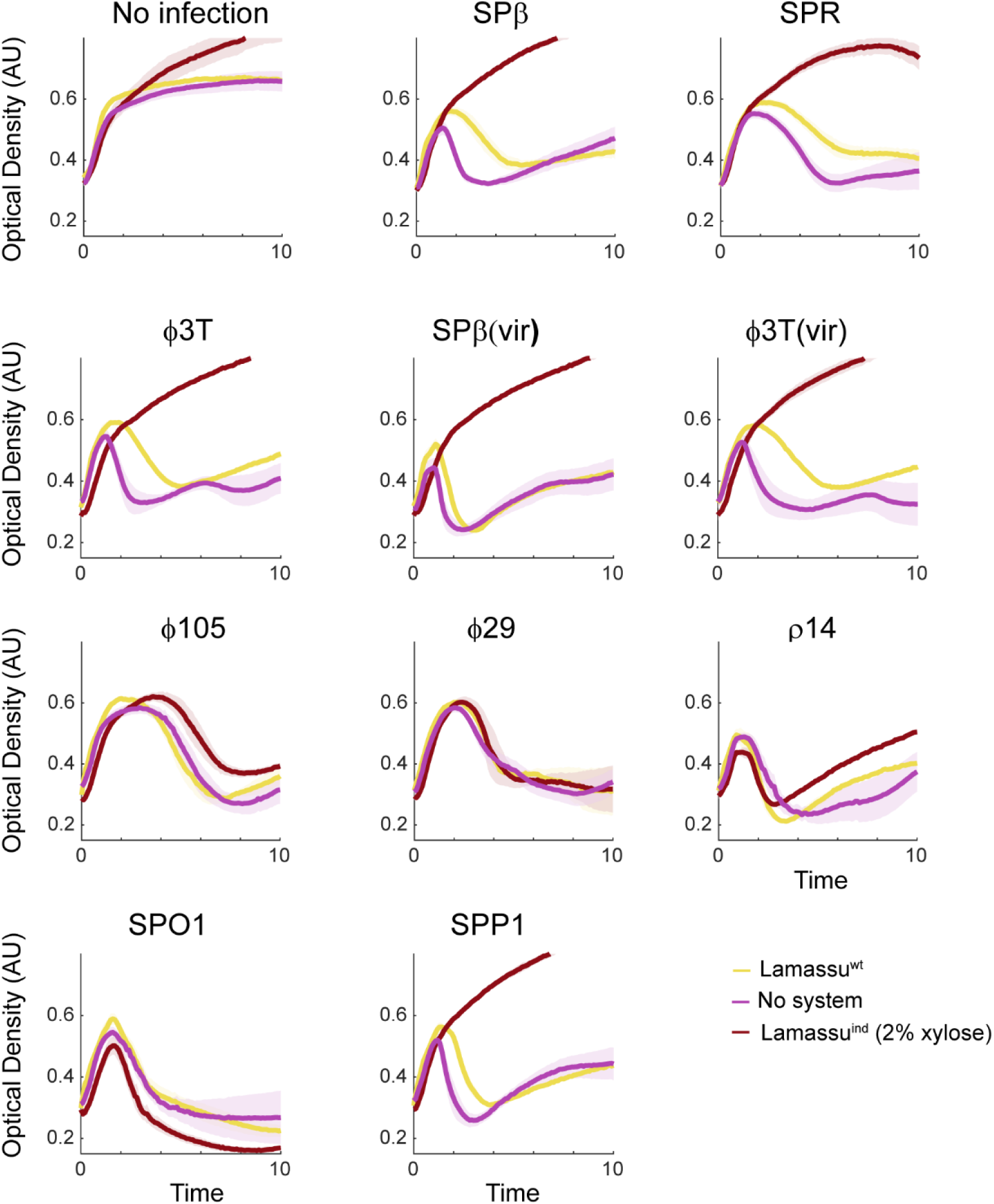
Growth curves of bacterial strains expressing Lamassu at different expression levels, infected by different phages. Mean OD of n=3 biological repeats and s.e.m (lighter shade) are shown.

**Supplementary Figure 6:**
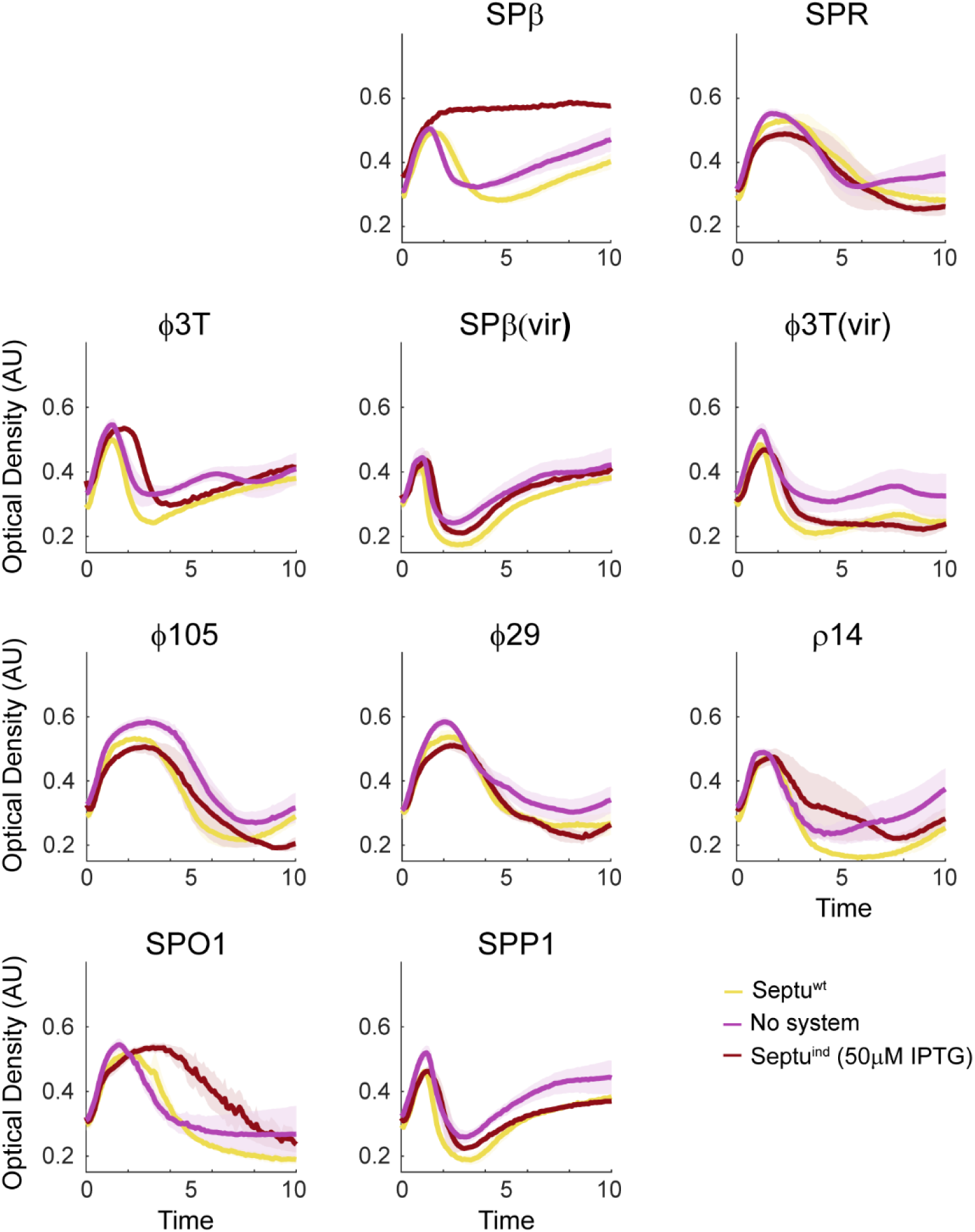
Growth curves of bacterial strains expressing Septu at different expression levels, infected by different phages. Mean OD of n=3 biological repeats and s.e.m (lighter shade) are shown.

**Supplementary Figure 7:**
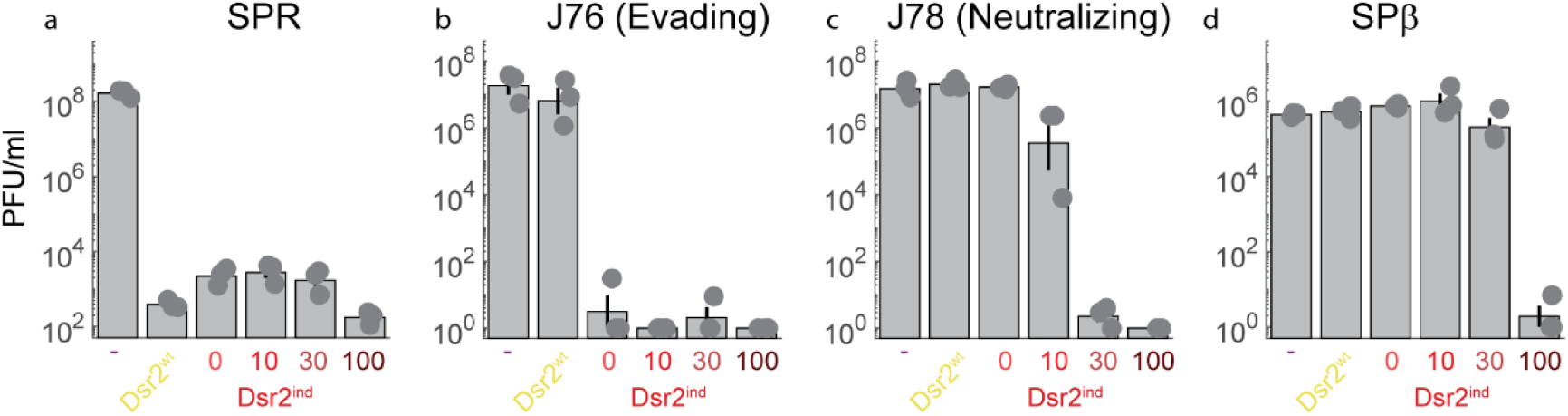
PFU/ml of EOP assays of different phages infecting strains expressing *dsr2* at different levels. Shown are geometric means and s.e.m of the log. Individual points represent n=3 biological repeats.

